# Nuclear ASC speck formation in microglia is associated with inflammasome priming and is exacerbated in LRRK2-G2019S Parkinson’s disease

**DOI:** 10.1101/2025.07.25.666809

**Authors:** Luca Ballotto, Thomas Baratta, Helena Winterberg, Cédric Dusanter, Sara Sambin, Jean Christophe Corvol, Ludovica Iovino, Luigi Bubacco, Olga Corti, Elisa Greggio, Salvatore Novello

**Author notes:** Corresponding author: Elisa Greggio. These authors contributed equally (Luca Ballotto, Thomas Baratta).

## Abstract

Neuroinflammation is increasingly recognized as a central pathological mechanism in Parkinson’s disease (PD), a progressive neurodegenerative disorder characterized by the selective loss of dopaminergic neurons and variety of motor and non-motor symptoms. The NOD-like receptor family pyrin domain-containing 3 (NLRP3) inflammasome and its adaptor protein ASC play a critical role in initiating and maintaining inflammatory responses in the central nervous system. Although its acute activation is beneficial for host defense and homeostasis, chronic activation of the inflammasome has been associated with the pathogenesis of PD. Another key contributor to neuroinflammation is the leucine-rich repeat kinase 2 (LRRK2), particularly the G2019S mutation associated with PD, which has been shown to exacerbate inflammatory signaling in microglia and peripheral immune cells. However, the interaction between LRRK2 and the NLRP3 inflammasome pathway remains poorly understood. In this study, we investigated the role of LRRK2-G2019S in the priming and activation dynamics of the NLRP3 inflammasome using mouse primary microglia and human monocyte-derived microglia-like cells (hMDMi). We observed that LRRK2-G2019S microglia exhibit increased expression of NLRP3 under basal conditions and spontaneous formation of ASC specks within the nucleus, an unexpected subcellular location not previously reported in microglia. Interestingly, nuclear ASC specks also formed in wild-type microglia and hMDMi after lipopolysaccharide (LPS) priming but only progressed to cytosolic ASC specks and interleukin-1β release after subsequent exposure to canonical NLRP3 activators. These findings suggest that nuclear ASC specks may represent a primed state of inflammasome activation and propose a novel cellular phenotype associated with LRRK2-G2019S. Altogether, our results reveal a new layer of inflammasome regulation in microglia and implicate LRRK2-G2019S in the promotion of a pro-inflammatory state, which may predispose to chronic neuroinflammation in PD. These findings advance our understanding of glial immune regulation and highlight potential therapeutic targets in PD.

## Background

Parkinson’s disease (PD) is an age-related neurodegenerative disease characterised by progressive impairment of motor functions and a wide variety of nonmotor symptoms. The main neuropathological hallmarks of the disease are the selective loss of dopaminergic neurons within the substantia nigra pars compacta (SNpc), the progressive prion-like spreading of α-synuclein, and chronic neuroinflammation (Poewe et al., 2017). The LRRK2-G2019S gain of function mutation is by far the most common genetic mutation among familial cases (Bouhouche et al., 2017; Paisán-Ruíz et al., 2004; Zimprich et al., 2004). Remarkably, while most PD cases are idiopathic with no clear genetic cause, LRRK2-associated PD patients exhibit clinical similarities to idiopathic PD patients (Rocha et al., 2022). Furthermore, increased LRRK2 activity has been observed in idiopathic PD, suggesting that the pathogenic mechanisms driving LRRK2-PD may also play a role in sporadic forms of the disease (Rocha et al., 2022).

Interestingly, preliminary single nucleus RNA sequencing data indicate that in the aging brain, and in idiopathic PD brains, LRRK2 levels are selectively up-regulated in microglia, suggesting that this cell type may be an important driver of dopaminergic degeneration (Duffy et al., 2023; Martirosyan et al., 2024). In fact, many studies have shown that LRRK2 mutations enhance the release of inflammatory cytokines in response to different types of stimuli, such as lipopolysaccharide (LPS) and α-synuclein, indicating that LRRK2-mediated neuroinflammation is a possible pathogenic mechanism in the onset and progression of PD (Pajarillo et al., 2023; Panagiotakopoulou et al., 2020; Russo et al., 2018).

IL-1β release has been found to be consistently increased by immune cells harbouring the G2019S mutation, and IL-1β levels are significantly higher in the blood and CSF of PD patients as well (Bessler et al., 1999; Mogi et al., 1996; Qin et al., 2016). The production of IL-1β depends on the activity of the NLR Family Pyrin Domain Containing 3 inflammasome (NLRP3), a highly-conserved multi-protein complex that orchestrates the release of inflammatory cytokines in response to a wide variety of pathogen- and damage-associated molecular patterns (PAMPs/DAMPs) (Kelley et al., 2019). Although the acute activation of the NLRP3 pathway is critical for the maintenance of brain homeostasis, its chronic activation is implicit in the pathology of PD, as the mRNA and protein levels of the inflammasome components are upregulated in the brain of Parkinson’s patients and in several preclinical models of PD (reviewed in Jewell et al., 2022). The activation mechanism of NLRP3 involves two steps (Figure 1A). First, the presence of PAMPs/DAMPs triggers NF-kB-dependent upregulation of the sensor protein NLRP3 and the pro-IL-1β which is known as inflammasome priming. Subsequently, a second stimulus, such as ATP or ROS, triggers NLRP3 oligomerization and the recruitment of the adaptor protein ASC, which is redistributed from the nucleus to a perinuclear aggregate known as the ASC speck (Bryan et al., 2009). ASC recruits the pro-caspase-1 zymogen through CARD-CARD (Caspase recruitment domains) interactions, which triggers dimerization and autoproteolytic activation of the effector protein, caspase-1. Finally, active caspase-1 cleaves pro-IL-1β and pro-IL-18 into their biologically active forms (Kelley et al., 2019).

**Figure 1.**
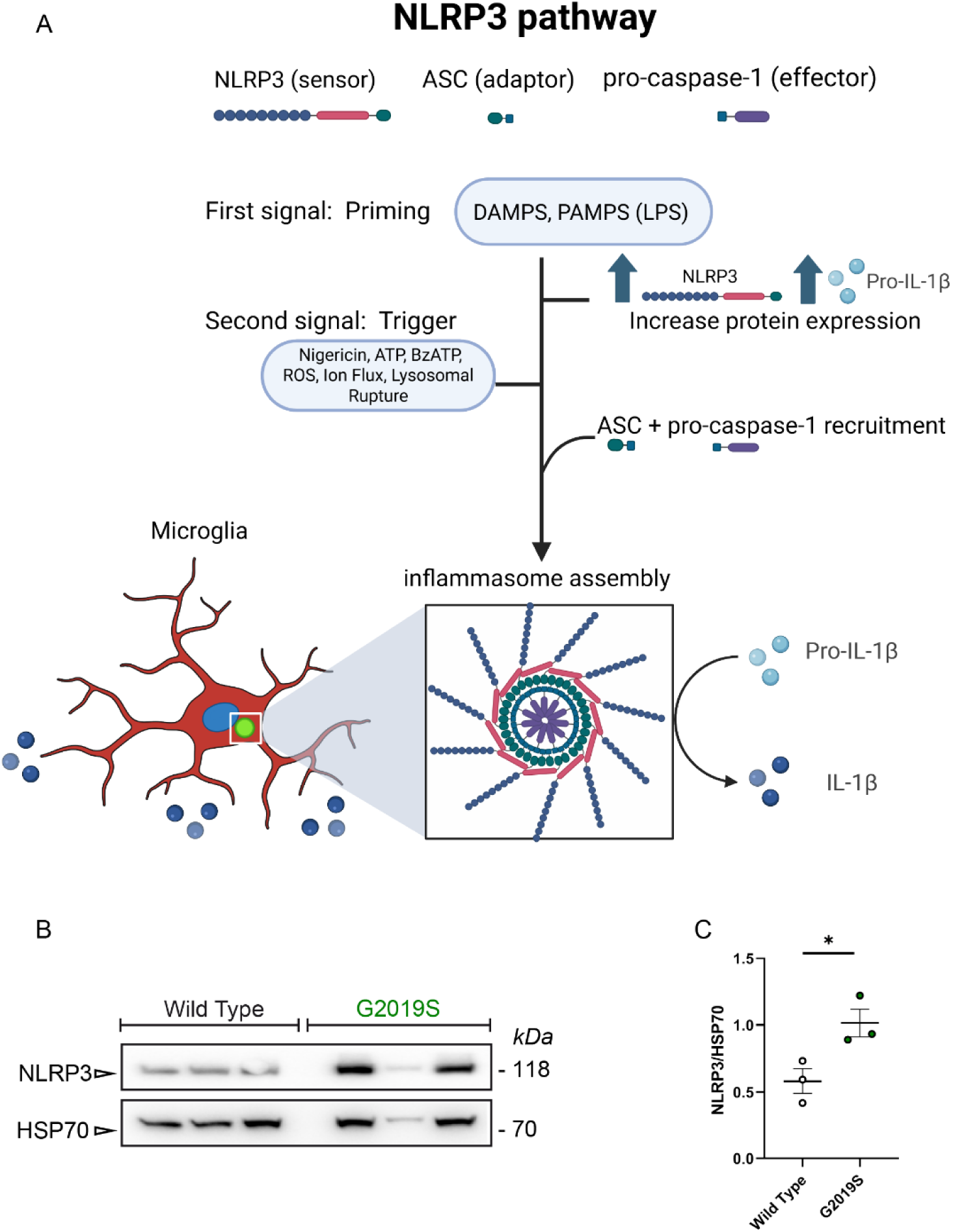
The NLRP3 protein is enriched in LRRK2 G2019S cortical microglia. (A) Schematic representation of the canonical NLRP3 inflammasome activation pathway. Created in BioRender. Ballotto, L. (2026) https://BioRender.com/1ozgh49.(B) Western blot from untreated primary Wild Type microglia and LRRK2 G2019S microglia. (C) Quantification of NLRP3 protein level normalized for the housekeeping HSP70. n=3 biological replicas for each genotype. The normality of the distribution was confirmed by Shapiro–Wilk test. *p < 0.05 by Student’s t-test. Results are expressed as mean ± SEM.

The gain-of-function LRRK2-G2019S mutation has been shown to exacerbate inflammatory responses in immune cells (Russo et al., 2022; Wallings et al., 2020). However, the detailed molecular mechanisms underlying this phenomenon are only partially understood. While LRRK2 has been robustly linked to NF-κB signalling (Moehle et al., 2012; Pajarillo et al., 2023; Russo et al., 2015), the molecular pathways and the potential connection between LRRK2 and inflammasome activation are still poorly resolved.

Here, we investigate the impact of the LRRK2 G2019S mutation on the NLRP3 inflammasome pathway in mouse primary microglia and human monocyte-derived microglia differentiated from LRRK2-PD patients. We report a novel phenotype characterized by intranuclear ASC speck formation in LPS-primed microglia, which is phenocopied by the G2019S mutation. Our results suggest that intranuclear ASC aggregation is a microglia-specific NLRP3 regulatory mechanism and that LRRK2 primes microglia for NLRP3 activation by preassembling inflammasome components.

## Materials and methods

### Mouse primary microglia cultures

Microglia cells were derived from postnatal days 0-4 Wild Type and *Lrrk2* G2019S knock-in homozygous mouse brains (C57BI/6J). Cerebral cortices were dissociated in cold PBS and separately resuspended in culture medium, which is composed by complete DMEM medium (Biowest^TM^, L0104) supplemented with 10% heat-inactivated FBS (Corning, 35-071-CV) and 1% penicillin and streptomycin (Biowest^TM^, L0022), so that each falcon tube contained 4 hemicortexes. The cell suspension was allowed to settle for 2 minutes, then the top fraction was collected and centrifuged at 200 rcf for 10 min. Cell pellets were resuspended in culture medium, seeded in poly-L-lysine (Sigma Aldrich^TM^, P4707) coated T75 cell culture flasks, and allowed to seed overnight. After 1 day, the medium was replaced with DMEM supplemented with 10% L929 conditioned medium, to increase microglia cell proliferation. At 10 days, microglia cells were mechanically detached from the mixed culture. The medium containing isolated microglia was collected, centrifuged at 200 rcf for 10 min, and resuspended in culture medium.

### L929-conditioned medium preparation

L929 is a fibroblast cell line that secretes Macrophage colony-stimulating factor (M-CSF), therefore, L929-conditioned medium can be used to promote microglia proliferation. L929 cells were seeded in a T175 flask and incubated at 37°C, 5% CO2 for 20 days. The L929 conditioned medium was collected on day 20 and filtered with sterile 0.22 μm pore size filters (Sarstedt^TM^, 83.3942.501). Finally, the filtered L929-conditioned medium was aliquoted and stored at -20°C until use.

### THP-1 cell culture

THP-1 cells stably expressing ASC-mCerulean or mCerulean were cultured in suspension at 37 ° C, with 5% CO_2_, in RPMI 1640 (Sigma Aldrich^TM^), supplemented with 10% heat-inactivated FBS (Corning, 35-071-CV), 1% Penicillin-Streptomycin solution (Biowest^TM^, L0022), 10mM HEPES (Euroclone, ECM0180D), 1mM sodium pyruvate (Sigma, S8636), 4.5g / L glucose (Sigma Aldrich^TM^, G7021) and 0.05 mM 2-Mercaptoethanol (Gibco^TM^, 21985023). 2-Mercaptoethanol was always added fresh the day of use. After seeding, THP-1 differentiation in macrophages was achieved by adding 25 ng/ml of phorbol 12-myristate 13-acetate (PMA) for 24h followed by washing and incubation for another 24h in fresh medium. THP-1 differentiated macrophages were treated with either LPS (100 ng/mL) + Nigericin or LPS alone (100 or 200 ng/mL) or vehicle (DMSO + Ethanol).

### Cellular treatment

Cells were seeded at a density of 2.5 x 10^5^ cells/well on poly-L-lysine-coated coverslips in a 24-well plate and allowed to adhere overnight at 37°C, 5% CO_2_. Before treatments, cells were starved overnight in FBS-free DMEM + penicillin/streptomycin and N-2 supplement (Gibco^TM^, 17502048). All cell treatments were performed in FBS-free cell culture medium. NLRP3 canonical activation was induced using 5µM Nigericin (Merck^TM^, 481990) for 30 minutes, following priming with 20 ng/mL LPS for 3,5 hours. After stimulation, cell culture medium was collected and stored at -20°C for immunoblotting experiments. Cells were rinsed in PBS and fixed with 4% paraformaldehyde for 20 min for subsequent immunolabeling.

### Cell lysis and protein quantification

For Western blot experiments, primary microglia cells were seeded at a density of 250000 cells per well in a 24-well plate and allowed to adhere overnight. Cell culture medium was removed, and cells were rinsed once with PBS. An appropriate volume of f RIPA buffer (20 mM Tris-HCl pH 7.5, 150 mM NaCl, 1 mM Na2EDTA, 1 mM EGTA, 1% NP-40, 1% Sodium deoxycholate, 2.5 mM 30 sodium pyrophosphate, 1 mM β-glycerophosphate, 1 mM Na_3_VO_4_, deionized water) enriched with 1X protease inhibitors cocktail (Sigma Aldrich^TM^, 11836170001), was added to each well, and lysis was performed using a cell scraper with the multi-well plate on ice. The lysates were collected and transferred in 1.5 mL Eppendorf tubes and kept on ice for 30 minutes. The samples were centrifuged at 20238 rcf for 30 min at 4°C. The pellet containing the remaining cell debris was discarded and 20 supernatants were collected in a new tube and stored at -80°C until use. Protein concentration quantification was performed using the Pierce® BCA Protein Assay Kit (Thermo Scientific^TM^, 23225) according to the manufacturer’s instructions. Protein samples were diluted with RIPA buffer to the desired concentration and an appropriate amount of SDS sample buffer 4X (SB 4X, 200 mM TRIS pH 6.8, 8% SDS, 400 mM DDT, 0,4% bromophenol blue, 40% glycerol) was added to each sample.

### Protein precipitation

Before immunoblotting, proteins in cell culture supernatants were concentrated according to the methanol/chloroform precipitation protocol, as previously described (Jakobs et al., 2013). Briefly, serum-free cell culture supernatants were collected from each well in 1.5 mL tubes (250 µL from two technical replicates were pooled) and centrifuged at 800 rcf for 5 min at room temperature to remove cell debris. 500 µL of methanol and 125 µL of chloroform were added to each tube. Samples were extensively vortexed and centrifuged at 13,000 rcf for 5 minutes, resulting in the formation of an upper aqueous methanol phase, a protein interphase, and a lower chloroform phase. The upper phase was carefully discarded without disturbing the interphase and 500 µL of methanol was added to the samples, which were vortexed and centrifuged as before. At this point, the supernatant was discarded and the protein pellet at the bottom of the tube was allowed to dry at 55°C. The pellet was resuspended in 40 µL of 1 x SDS sample buffer. The samples were vortexed and boiled at 95°C for 5 minutes.

### Immunoblotting

Soluble proteins from cell lysates were fractionated by SDS-PAGE using a pre-casted gel with a gradient concentration of 4-20% of acrylamide (GenScript^TM^, M42015), in combination with Tris-MOPS running buffer (Genscript^TM^, M00138). Polyacrylamide gels were subjected to electrophoresis at 130V for 1.5 hours. Proteins were transferred to polyvinylidene fluoride (PVDF) membranes under semi-dry conditions using the Trans-Blot® TurboTM transfer system (Bio-Rad^TM^, 1704150) with the 1X Transfer Buffer (5X Bio-Rad transfer buffer 10026938, 20% ethanol, and deionized water), at 25V for 20 minutes (high molecular weight protocol). Molecular weight markers (Bio-Rad^TM^, 1610374) were loaded in at least one lane for each gel to infer the molecular weight of the sample proteins. The precipitated proteins of the cell culture medium were blotted as previously described with slight modifications: SDS-PAGE was performed using a 15% acrylamide gel and Tris-Glycine running buffer for 2 hours at 100 V and proteins were transferred to Nitrocellulose membranes. PVDF and nitrocellulose membranes were incubated for one hour in agitation in blocking solution (5% skim milk in Tris-Tween Buffered Saline, TBS-T) and overnight with the primary antibody diluted in blocking solution at the appropriate concentration (Table 1). The membranes were washed three times in Tris buffered saline with 0.1% Tween® 20 detergent (TBS-T) and incubated with appropriate HRP-conjugated secondary antibody for 1 hour (Table 1). The membranes were washed three more times to remove excess antibodies and Immobilon® Forte Western HRP Substrate (Millipore^TM^, WBLUF0500) was applied to the membranes for a couple of minutes. HRP signal was visualized using VWR® Imager Chemi Premium.

**Table 1.** Antibodies and dilutions.

| Antibody | Source | Identifier | Host | Dilution |
| --- | --- | --- | --- | --- |
| ASC | Adipogen | AG-25B-0006-C100 | Rabbit | 1:100 |
| Caspase-1 p20 | Genetech | 4B4.2.1 | Rat | 1:100 |
| CD11b | Abcam | ab1211 | Mouse | 1:200 |
| NLRP3 | Adipogen | AG-20B-0014-C100 | Mouse | 1:300 |
| Goat anti-rabbit<br>Alexa-Fluor 488 | Abcam | ab150077 | Goat | 1:200 |
| Goat anti-mouse<br>Alexa-Fluor 568 | Abcam | ab175473 | Goat | 1:200 |

### Immunostaining and epifluorescence imaging

Cells were permeabilized with 0.1% Triton X-100 in PBS for 20 min and saturated with blocking buffer containing 5% FBS in PBS for 1 h at RT. Primary antibodies (Table 1) diluted in blocking solution were incubated overnight, in a humidified chamber at 4°C. The next day, the coverslips were rinsed in PBS three times and incubated for 1 h at RT with secondary antibodies in blocking solution (Table 1). After repeated washes, cells were incubated for 5 min with Hoechst 33258 (Invitrogen^TM^, 1:10000 in PBS) to visualize nuclei. Coverslips were then mounted using Mowiol (Calbiochem^TM^, 475904) and allowed to dry overnight at 4°C. From each experimental condition, five images were acquired with a Leica DMI4000 epifluorescence microscope at a magnification of 40X. Cells positive for nuclear and cytosolic ASC specks were separately counted using the ImageJ software cell counter plugin and normalized for the number of nuclei. The total number of cells analyzed for each condition were: 892 Wild Type vehicle, 672 Wild Type LPS + Nigericin, 659 LRRK2 G2019S vehicle, 538 LRRK2 G2019S LPS + Nigericin.

### Statistical analysis

Statistical analyses were performed using GraphPad Prism^TM^ 8.0.2 software. All quantitative data are expressed as mean ± SEM (standard error of the mean). The appropriate statistical test was selected after using the Shapiro-Wilk test to evaluate the normality of the data. After confirming the Gaussian distribution, the unpaired parametric student’s t-test was carried out to evaluate the statistical significance between two groups in the western blot and immunocytochemistry experiments. To compare the effects of genotype and multiple treatments in the LPS priming immunocytochemistry experiments, Two-way ANOVA was used, followed by the post-hoc Šídák’s multiple comparisons test.

### Human monocyte-derived microglia-like cells (hMDMi) cultures

Blood samples were collected within the Clinical Investigation Center for Neurosciences at the Pitié-Salpêtrière Hospital (Paris, France) in PD patients included in the NS-PARK cohort (sponsored by Inserm, NCT04888364 (clinicaltrials.gov)) and in healthy control individuals participating in the BIOMOV cohort (sponsored by APHP, NCT05034172 (clinicaltrials.gov)). Both studies were conducted in accordance with Good Clinical Practice, received approval from an ethical committee and regulatory authorities according to local regulations. Written informed consent was obtained from all subjects undergoing sample collection. Preliminary experiments were performed on MDMis from healthy donors: these subjects were not affected by any chronic or acute inflammatory disease, and they did not suffer from any neurological disorder. A case-control study was then performed on MDMis from *LRRK2*-(G2019S mutation) patients and matched controls (Table 2).

**Table 2.** List of subjects included in the case-control study.

| Status | Age | Withdrawal date | Sex | Mutant gene | Mutation | Onset of symptoms | Disease duration |
| --- | --- | --- | --- | --- | --- | --- | --- |
| control | 60 | 23/05/2022 | F | NA | NA | NA | NA |
| patient | 57 | 23/05/2022 | F | LRRK2 | G2019S | 2018 | 4 yrs |
| control | 57 | 31/08/2022 | M | NA | NA | NA | NA |
| patient | 55 | 31/08/2022 | M | LRRK2 | G2019S | 2016-2018 | 4-6 yrs |
| control | 56 | 01/09/2022 | M | NA | NA | NA | NA |
| patient | 53 | 01/09/2022 | M | LRRK2 | G2019S | 2014 | 8 yrs |
| control | 53 | 16/09/2022 | M | NA | NA | NA | NA |
| patient | 53 | 16/09/2022 | M | LRRK2 | G2019S | n.d. | n.d. |

Peripheral blood mononuclear cells (PBMCs) were isolated from buffy coats following density gradient centrifugation in Ficoll (Sigma Aldrich^TM^, Histopaque, 10771), based on an established protocol (Sellgren et al., 2017). Cell number was determined using the automated cell counter ADAM™-MC. PBMCs were frozen until usage at -80°C in heat-inactivated FBS (Gibco^TM^, A3840401) supplemented with 10% dimethyl sulfoxide (DMSO, Sigma^TM^, D8418). Cryovials were thawed in water bath at 37°C. The cellular suspension was centrifuged at 300g for 5 min in RPMI Glutamax 1640 (Gibco^TM^, 61870044) enriched with 1% Penicillin/Streptomycin (P/S, Gibco^TM^, 15140122) and 10% FBS. Cell pellets were resuspended in the same medium. PBMCs were counted again and then seeded onto Geltrex (Gibco^TM^, A3840401)-coated 8-well chamber slides (ibidi^TM^ 8 Well Chamber, 80826), at a density of 8.4 x 10^5^ cells/cm^2^. After overnight incubation, performed under standard humidified culture conditions (37°C, 5% CO_2_), the culture medium was replaced with a serum-free medium. To induce MDMi differentiation, monocytes were incubated up to 14 days with human recombinant cytokines IL-34 (100 ng/ml; R&D Systems^TM^, 5265-IL-010/CF) and GM-CSF (10 ng/ml; R&D Systems^TM^, 215-GM-050/CF), as previously described (Sellgren et al., 2017).

### hMDMi treatments and medium analysis

During treatment, cells were cultured in cell culture medium supplemented with 1% FBS. MDMi were treated with 10 ng/ml LPS (Sigma Aldrich^TM^, 0111:B4) for 3.5h as a priming stimulus. During the last 30 min, 300μM BzATP (Sigma Aldrich^TM^, B6396) was added to the medium to trigger NLRP3 activation. After stimulation, cells were fixed using 4% paraformaldehyde (PFA) for 14 min; cell culture medium was collected and stored at -20°C for subsequent enzyme-linked immunosorbent assay (ELISA). The medium was specifically analysed for the presence of the pro-inflammatory cytokines IL-1β (Human IL-1β/IL-1F2 DuoSet; R&D Systems^TM^, DY201-05).

### Immunostaining and confocal imaging

Cells were incubated for 1 h at RT in PBS-Triton 0.3% (Sigma Aldrich^TM^, T8787), enriched with 0.3 M Glycine (Sigma Aldrich^TM^, G8898) and 5% BSA (SigmaTM, A3294). Primary antibodies ASC (1:300; AdipoGen^TM^, AG-25B-0006-C100) and IBA-1 (1:5000; Synaptic Systems^TM^, 234009) were diluted in PBS-BSA 5%-Tween 0.05% (Sigma Aldrich^TM^, P7949). Incubation with primary antibodies was performed overnight at 4°C in a humidified chamber. The day after, the wells were washed three times with PBS-Tween 0.05% and incubated for 50 min at RT with secondary antibodies Alexa-Fluor 488 and 568 (1:1000) in incubating solution. After repeated washes, cells were incubated for 10 min with DAPI (Abcam™, ab228549; 1:1000 in PBS-Tween 0.05%) to visualize nuclei. Slides were mounted using Fluoromount-G™ mounting medium (Invitrogen^TM^, 00-4958-02) and allowed to dry overnight at 4°C. Image acquisition was performed using an A1R-HD25 Nikon inverted confocal microscope. For each experimental condition, four images were acquired at 40X magnification. Quantification of cells positive for nuclear and cytosolic ASC specks was performed using the ImageJ software cell counter plugin and normalized for the number of nuclei. The total number of cells analyzed for each condition were: 1206 Wild Type vehicle, 1429 Wild Type LPS, Wild Type 1384 LPS + BzATP; 1382 LRRK2 G2019S vehicle, 1725 LRRK2 G2019S LPS, 964 LRRK2 G2019S LPS + BzATP.

## Results

### 1. NLRP3 levels are increased in LRRK2 G2019S cortical microglia

A previous study from our group pointed out LRRK2 as a positive regulator of nuclear factor kappa-B (NF-κB) signalling in microglia(Russo et al., 2019, 2018, 2015). Given that the hyperactive kinase activity of G2019S-LRRK2 could exacerbate NF-κβ activation and the NLRP3 coding gene is a target of NF-κβ, we evaluated NLRP3 protein levels in untreated cortical primary microglial cells derived from LRRK2-G0219S KI mice or littermate controls. Under unstimulated conditions, NLRP3 levels were significantly higher in G2019S microglia lysates compared to Wild Type microglia (Figure 1B,C), which could indicate that LRRK2 mutations primed microglia for inflammasome activation.

### 2. Nuclear ASC specks spontaneously form in primary LRRK2-G2019S cortical microglia

To further investigate the assembly of the NLRP3 inflammasome scaffold in primary microglia, we performed immunocytochemistry experiments to visualize the adaptor protein ASC (Figure 2). Monomeric ASC localizes in the cell cytoplasm and within the nuclei of macrophages in resting conditions (Bryan et al., 2009). Conversely, it oligomerizes into a large speck (around 1 μm diameter) when NLRP3 inflammasome scaffold assembly occurs, thus providing a readout for inflammasome assembly upstream of caspase-1 activation and IL-1β processing (Stutz et al., 2013). Given the higher steady state levels of NLRP3 in LRRK2-G2019S primary microglia, we set out to measure the number of ASC specks in LRRK2-G2019S microglia to evaluate the degree of inflammasome assembly in resting conditions. Primary microglia cells were stimulated with either LPS for 3.5 hours followed by the addition of Nigericin for the last 30 minutes or vehicle control. LPS acts as a priming signal and is required to increase the transcription of NLRP3 and pro-IL-1β, while Nigericin is a microbial pore-forming toxin that triggers potassium efflux and full inflammasome activation (Kelley et al., 2019). During image acquisition, we observed that ASC specks displayed distinct subcellular localization under specific experimental conditions (figure 2A). To quantify this, we performed confocal imaging, distinguishing between nuclear and cytosolic ASC specks. These were counted separately and normalized to the number of Hoechst-positive nuclei. Canonical inflammasome activation by established priming and activation signals (i.e. LPS and Nigericin), resulted in increased oligomerization of the ASC protein, which was predominantly located in the cytosolic or perinuclear area, regardless of genotype. Cytosolic specks displayed an increased trend in treated LRRK2-G2019S microglia, although not statistically significant (p=0,0828, unpaired t-test). Vehicle-treated Wild Type microglia display diffused ASC signal within the cells, preferentially in the nuclear area, with little to no ASC speck formation. On the other hand, vehicle-treated LRRK2-G2019S cortical microglia displayed increased formation of ASC specks, which were located exclusively within the nucleus (Figure 2C-D, Additional file 1). As expected, stimulation with LPS and Nigericin was associated with caspase-1 activation (Figure 2B).

**Figure 2.**
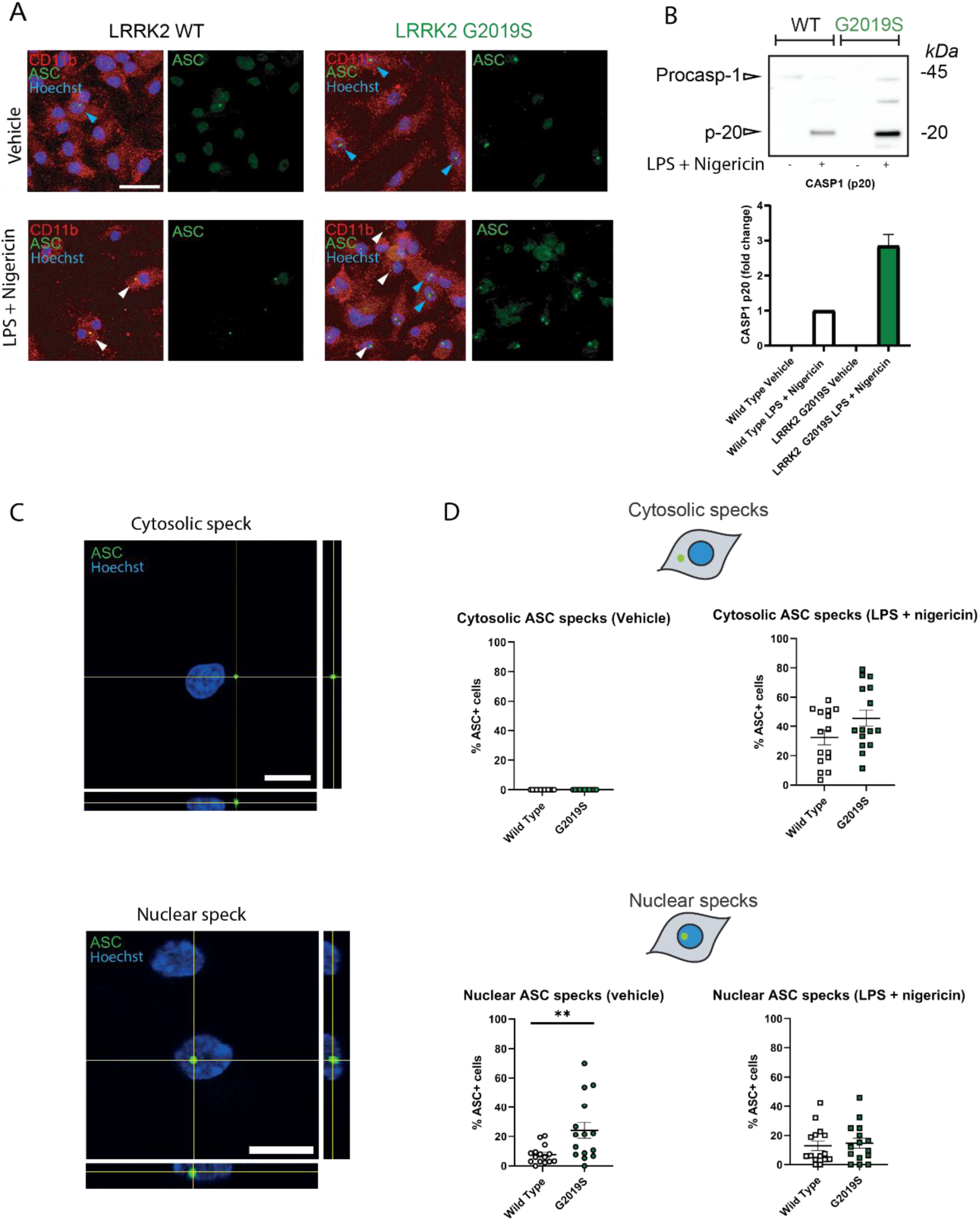
ASC speck formation in primary LRRK2-G2019S cortical microglia. (A) LRRK2-Wild Type and LRRK2-G2019S primary cortical microglia were exposed to either LPS + Nigericin or vehicle and stained with the anti-ASC antibody (green) and the anti-CD11b antibodies (red) Scale bar: 50 um, zoom 25 um. (B) Concentrated proteins from the cell culture medium of primary microglia were immunoblotted for active caspase-1 p20 fragment. Representative blot from n=2 independent experiments. Caspase-1 p20 signal was normalized on Wild Type LPS + Nigericin level. (C) The subcellular localization of ASC could be observed either in the cytosolic or nuclear area and was quantified separately. Scale bar: 10 um. (D) Quantification of nuclear or cytosolic ASC specks identified in basal conditions or after stimulation with LPS + Nigericin. Each data point represents to the average of 106-375 cells per field, from n=15 fields collected across 3 independent experiments. The normality of the distribution was confirmed by Shapiro–Wilk test. ** p <0.01 by Student’s t-test. Results are expressed as mean ± SEM.

### 3. LPS priming is sufficient to induce nuclear ASC speck formation in primary microglia

The detection of nuclear ASC aggregation in microglia is, to our knowledge, a novel finding that warrants further investigation, as it may be indicative of an intermediate step in the NLRP3 activation pathway. Since LRRK2-G2019S microglia also displayed higher levels of NLRP3 protein, we hypothesized that the nuclear formation of ASC specks may be related to inflammasome priming. To test this hypothesis, we treated primary microglia cells with either LPS + Nigericin (priming + trigger) or LPS alone, to assess the effect of inflammasome priming on ASC aggregation in microglia. Remarkably, Wild Type microglia, when primed with LPS for 3.5 hours, exhibited ASC speck formation exclusively within the nucleus (Figure 3A-B). Similarly, LPS-primed G2019S microglia also exhibited intranuclear ASC specks, but without significant difference compared to vehicle condition. On the contrary, perinuclear or cytosolic ASC specks were only detected when Nigericin activation was performed following priming, and, once again, a trend of increased cytosolic ASC specks was observed in LRRK2 G2019S microglia. Thus, inflammasome priming and activation have distinct effects on ASC speck formation and subcellular localization: cytosolic ASC specks are detected only in the presence of priming and trigger, whereas inflammasome priming alone is sufficient to induce nuclear ASC speck formation in microglia. These results suggest that the LRRK2-G2019S mutation primes microglia for inflammasome activation in a similar way as the LPS treatment does for Wild Type microglia. Interestingly, THP-1-differentiated macrophages only exhibit cytosolic or perinuclear ASC specks when primed with LPS or treated LPS + Nigericin, indicating that NLRP3 regulatory mechanisms may be different between microglia and macrophages (Supplementary Figure 1).

**Figure 3.**
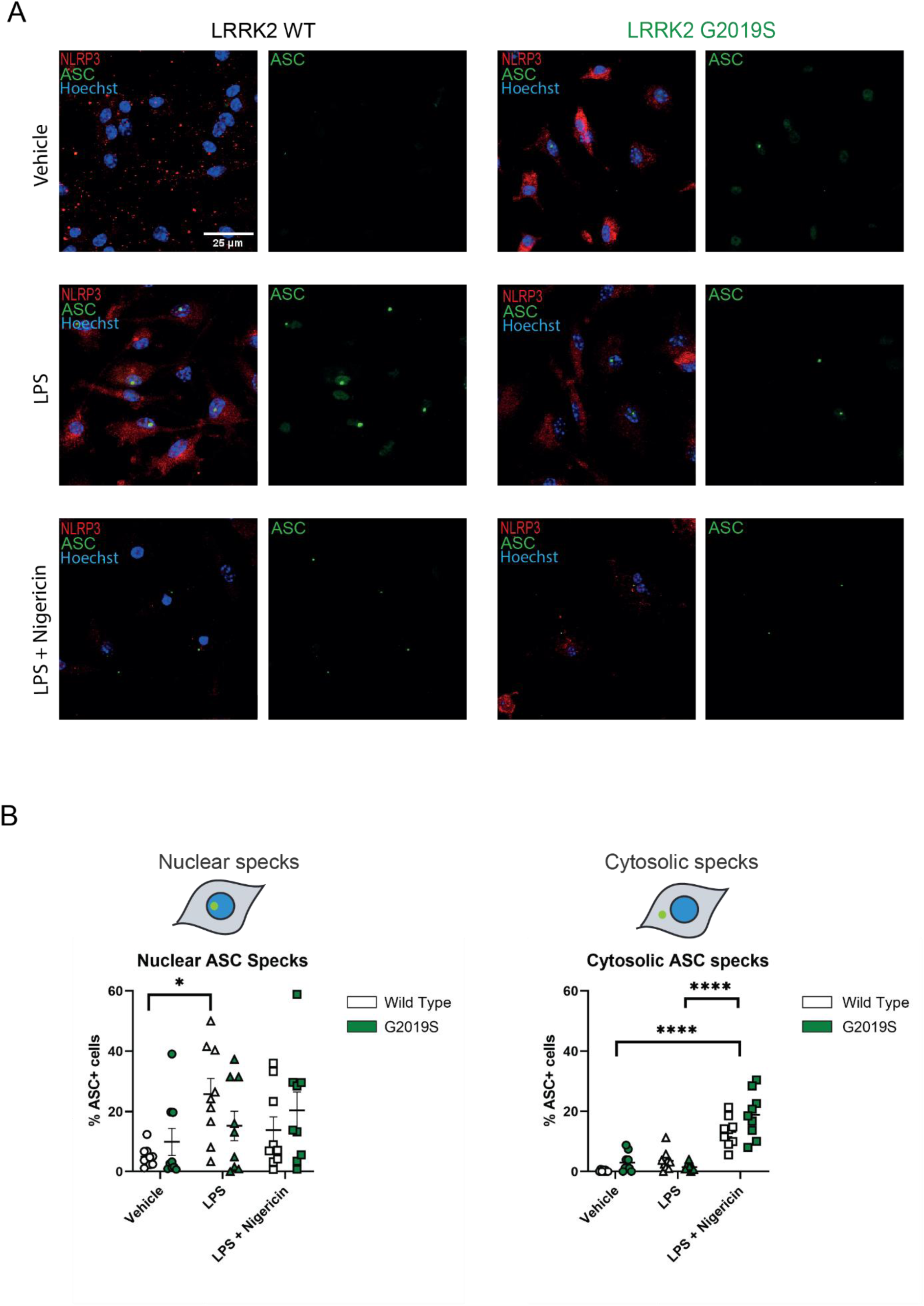
LPS priming induces nuclear ASC speck formation in primary microglia: (A) Representative acquisitions of primary cortical microglia from LRRK2-Wild Type and LRRK2-G2019S mice treated with LPS alone, LPS + Nigericin, or vehicle. (B) Quantification of nuclear and cytosolic ASC specks normalized on Hoechst positive nuclei. Each data point represents to the average of 27-233 cells per field, from n=9 fields collected across 3 independent experiments. The normality of the distribution was confirmed by Shapiro– Wilk test. * p <0.05, **** p <0.0001 by Two-Way ANOVA. The results are expressed as mean ± SEM.

### 4. Nuclear ASC specks assemble in PD patient-derived hMDMi

Nuclear ASC oligomerization is enhanced in primary cortical LRRK2-G2019S mouse microglia. If confirmed in human models, the accumulation of ASC specks within cell nuclei under basal conditions may represent a hallmark of LRRK2-PD, indicating the involvement of LRRK2 in the complex molecular pathway of NLRP3 inflammasome activation. To test this, we evaluated ASC oligomerization in human monocyte-derived microglia-like cells (hMDMis) from four LRRK2 PD patients in a case-control study (see Table 1). Following differentiation, hMDMis from both genotypes were divided into three groups: vehicle, LPS-treated and LPS+BzATP-treated. BzATP is an inflammasome trigger robustly validated in human microglia and microglia-like cells, including hMDMis (Heavener et al., 2024; Janks et al., 2018). In agreement with mouse microglia, ASC immunofluorescence revealed the presence of nuclear specks in absence of treatment and in a greater proportion of cells in the LRRK2-G2019S context compared to Wild-Type (Figure 4; p=0.0086). The proportion of cells positive for nuclear ASC specks significantly increased following priming with LPS in Wild-type cells; however, mutant cells did not show a further increase in nuclear ASC oligomerization, which remained lower than in Wild-type cells. Similar to inflammasome-triggered mouse microglia, BzATP exposure led to a decreased proportion of cells exhibiting nuclear ASC specks, and to a corresponding increase in the proportion of cells with cytosolic specks. Interestingly, we observed a stronger response to BzATP in LRRK2-G2019S cells compared to Wild Type (Figure 4; p<0.0001).

**Figure 4:**
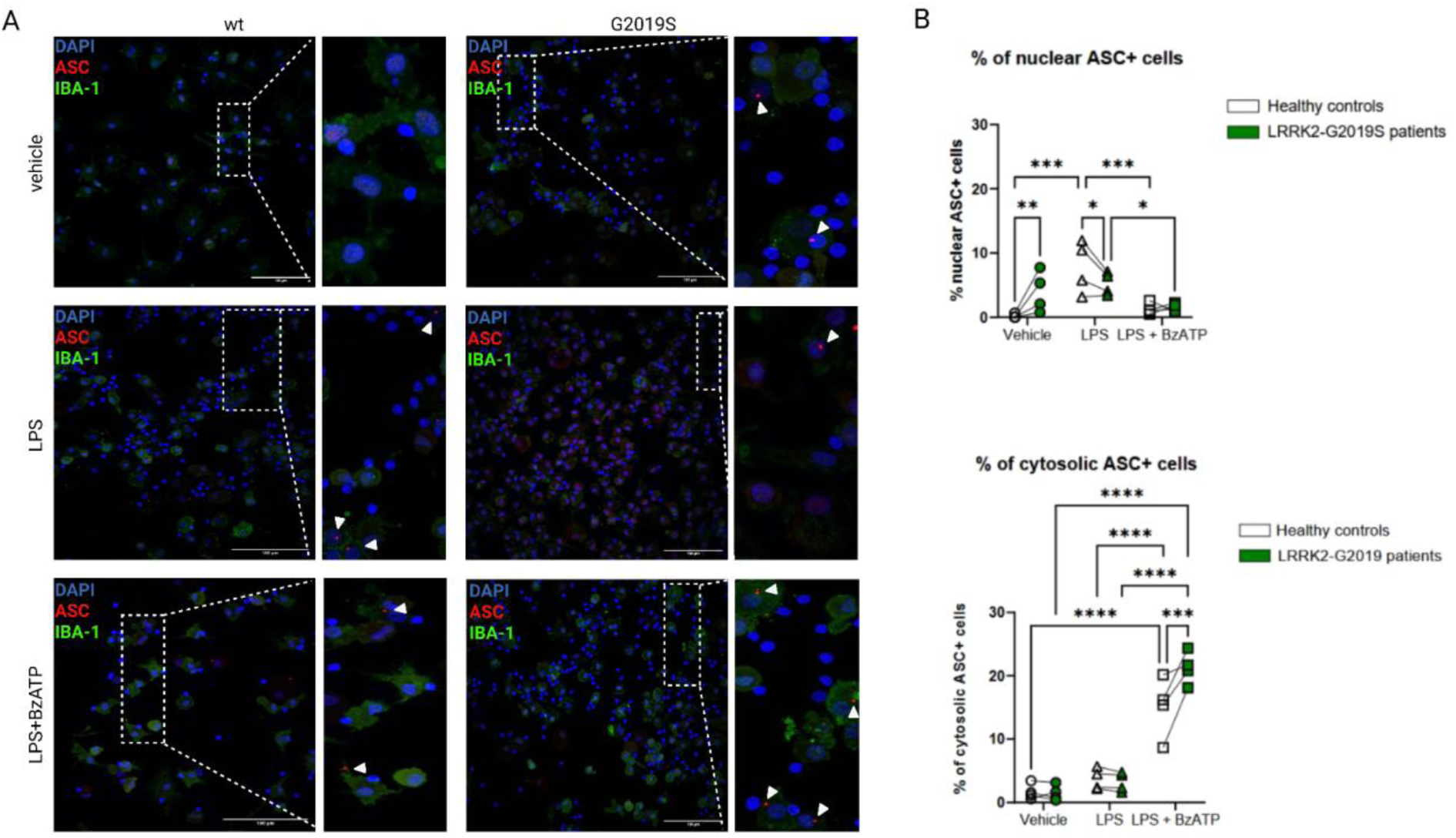
ASC speck formation in monocyte-derived microglia (MDMis) from LRRK2-G2019S PD patients and matched controls. (A) Representative images illustrating ASC (red) and IBA-1 (green) immunostaining. Scale bar x = 100 µm. (B) Quantification of nuclear and cytosolic ASC specks normalized on DAPI positive nuclei. Each point in the graph corresponds to the mean value for each individual, calculated considering 16 - 261 cells per field, from n = 4 fields per individual. The normality of the distribution was confirmed by Shapiro–Wilk test. * p <0.05, **** p <0.0001 by Two-Way ANOVA. The results are expressed as mean ± SEM.

In hMDMis from three different healthy donors, we observed that the relative number of cells presenting nuclear ASC specks significantly increased in response to LPS priming (Figure 5B, p < 0.0001). Within the LPS-treated group, the proportion of cells positive for nuclear ASC specks was higher than the proportion of cells with cytosolic ASC specks (Figure 5B; p<0.001). This result further strengthens the hypothesis that ASC oligomerization occurs within the cell nucleus as a consequence of priming.

**Figure 5:**
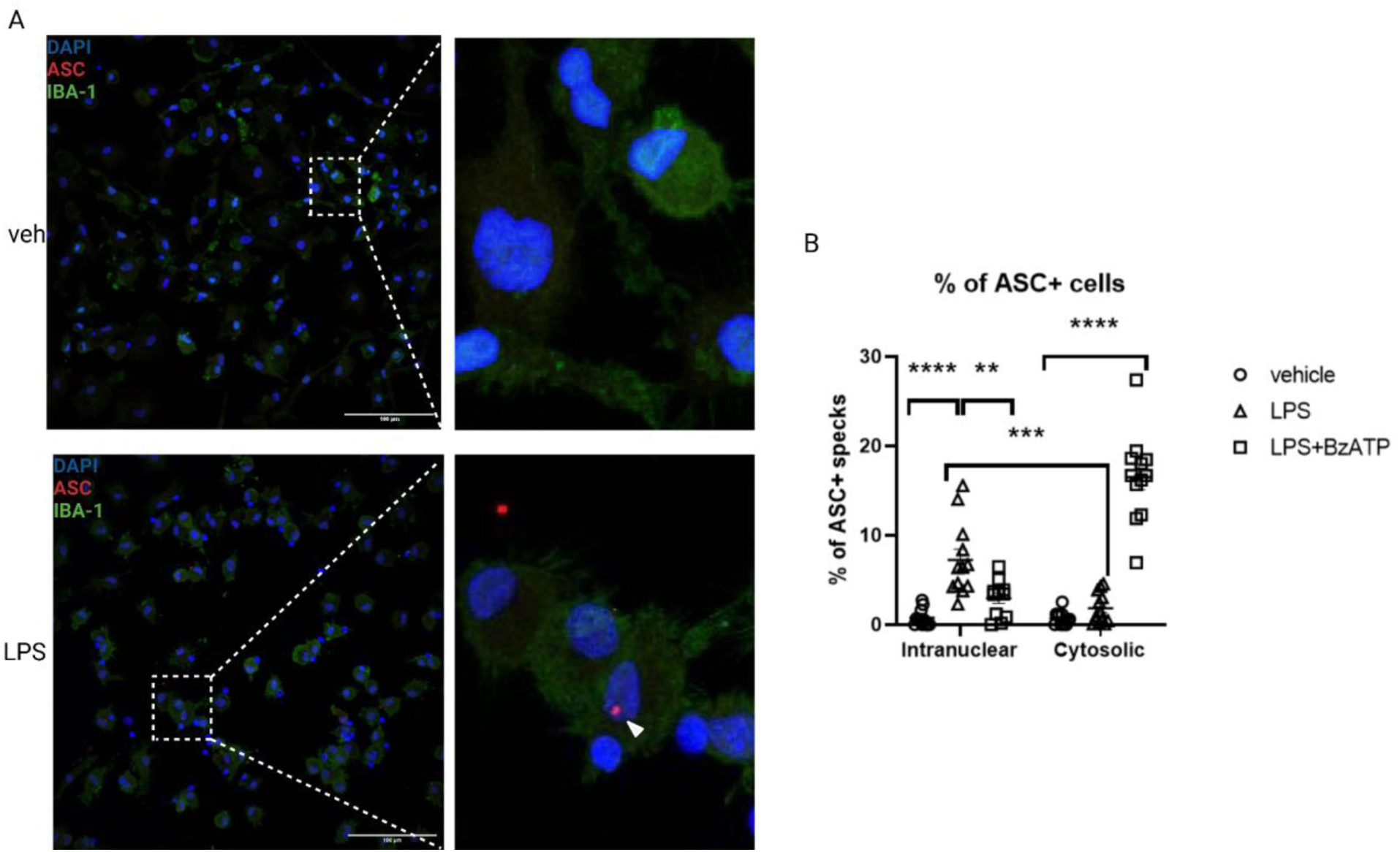
Nuclear ASC specks in resting LRRK-G2019S microglia. (A) Representative images illustrating ASC (red) and IBA-1 (green) immunostaining. Scale bar x = 100 µm. (B) Quantification of nuclear and cytosolic ASC specks normalized on DAPI positive nuclei. Each point in the graph corresponds to the average of 45 - 276 cells per field, from n = 12 fields collected across 3 independent experiments. The normality of the distribution was confirmed by Shapiro–Wilk test. * p <0.05, **** p <0.0001 by Two-Way ANOVA. The results are expressed as mean ± SEM.

### 5. Increased IL-1β release in PD patient-derived hMDMi

The NLRP3 inflammasome regulates the neuroinflammatory response through the release of pro-inflammatory cytokines, such as IL-1β and IL-18. We performed an ELISA assay on the stimulation medium following hMDMis treatment, to quantify IL-1β release in each experimental condition. This parameter provides direct and specific evidence of the complete NLRP3 activation.

As illustrated in Figure 6, LRRK2-G2019S cells released a higher amount of IL-1β in response to LPS+BzATP treatment, indicating overactivation of the NLRP3 inflammasome pathway consistent with the corresponding increase in cytosolic ASC specks. As expected, no IL-1β release was detected under the other conditions.

**Figure 6:**
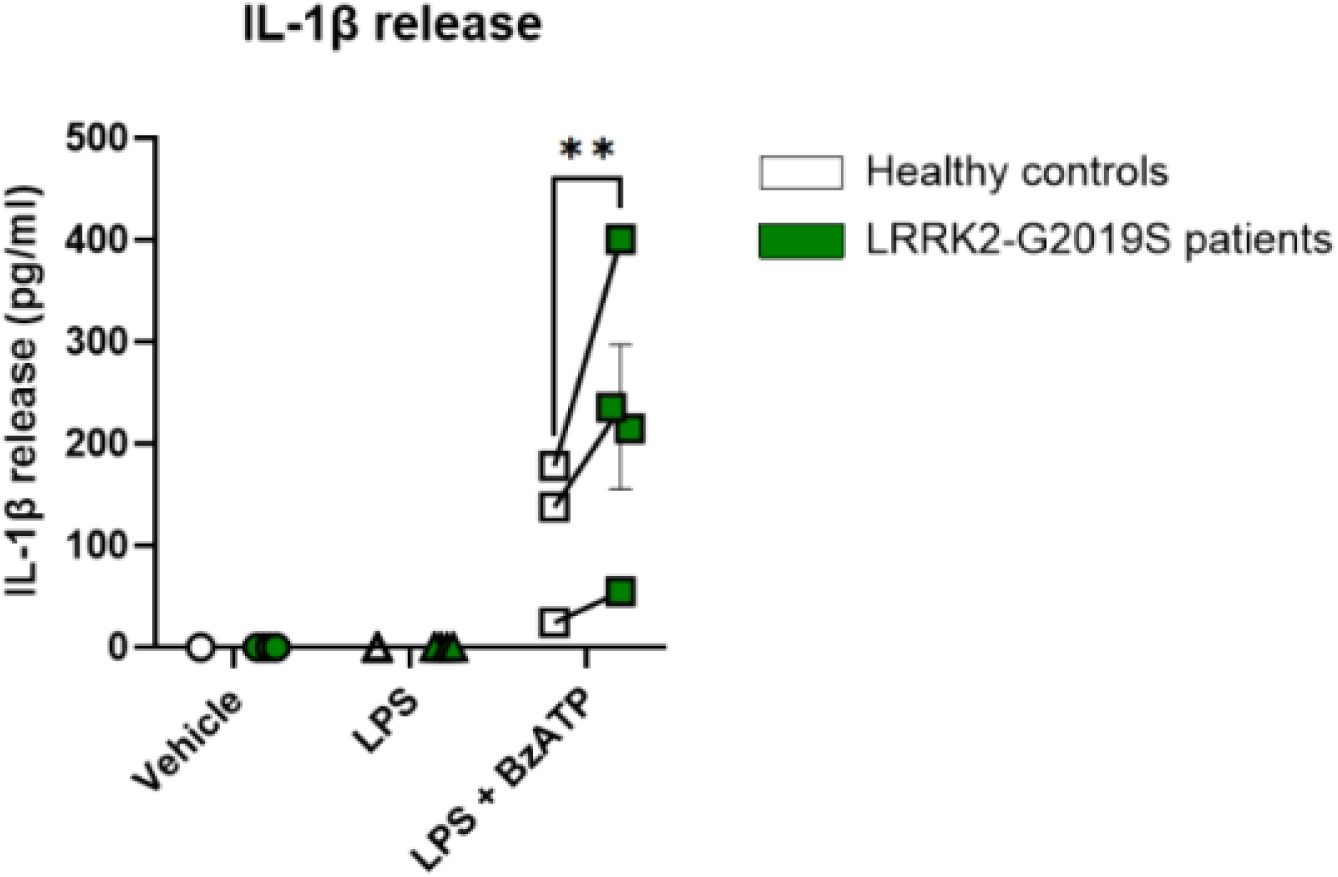
Quantification of IL-1ꞵ release in hMDMis from LRRK2 G2019S patients and matched controls. Each point in the graph represents the mean value for each individual, calculated based on n = 3 independent replicates. The normality of the distribution was confirmed by Shapiro–Wilk test. * p <0.05, **** p <0.0001 by Two-Way ANOVA. The results are expressed as mean ± SEM.

## Discussion

The results of this study suggest a role for the PD-linked kinase LRRK2 in the priming of microglia for NLRP3 inflammasome activation. The observed up-regulation of NLRP3 in LRRK2-G2019S primary microglia, coupled with the increased release of IL-1β from patient-derived HMDMi, enforces the growing body of evidence pointing to LRRK2-mediated neuroinflammation as a critical mechanism in the pathogenesis of PD.

Regulatory mechanisms of inflammasome scaffold assembly and ASC protein redistribution are poorly investigated, especially in the context of microglia. One study reported that human macrophages in resting condition display increased levels of non-aggregated ASC within the nucleus compared to the cytosol, suggesting that nuclear ASC localization could serve as a regulatory mechanism that prevents its interaction with NLRP3 in the cell cytosol in resting cells (Bryan et al., 2009). Similarly, Wild Type untreated microglia from our study displayed a diffused ASC signal throughout the cell, which peaked in the nuclear region, while in the resting mutant LRRK2-G2019S microglia, ASC specks could be detected within the nucleus. Importantly, a single perinuclear or cytosolic ASC speck is visible in cells treated with LPS + Nigericin, while up to two ASC specks could be frequently detected within a single nucleus in vehicle-treated LRRK2-G2019S cortical microglia. Another difference between intranuclear and cytosolic ASC specks is that diffused ASC signal can be detected in proximity to the intranuclear speck, but not to the cytosolic speck (Figure 3). This could indicate that monomeric ASC begins aggregating first in the nucleus of microglia, forming the nuclear ASC speck during inflammasome priming. Then, once the speck is relocated to the cytosol, all remaining monomeric ASC is recruited to the cytosolic speck and complete inflammasome activation is achieved.

The presence of intranuclear ASC specks has been previously observed in other cell types, but the functional implication of this peculiar localization is not fully understood and may be cell-type specific. The first report of intranuclear ASC specks was in a study on ASC speck formation dynamics in HeLa cells from 2010 (Cheng et al., 2010). Intranuclear ASC speck formation in HeLa cells first depleted the diffused nuclear ASC signal, followed by a slower depletion of cytoplasmic diffused ASC (Cheng et al., 2010). In contrast, nuclear ASC specks in microglia from our study are present in concomitance with diffused ASC signal, indicating a cell-type specific difference in ASC speck formation dynamics. Unfortunately, the functional implications compared with the classical perinuclear location of the ASC specks were not further discussed in the study. A second report of nuclear ASC specks comes from a study of HMVEC cells infected with nuclear replicating herpes virus (Kerur et al., 2011). Infected HMVEC cells displayed a speckled pattern of ASC colocalizing with caspase-1 in the nucleus after 2 hours, and a redistribution of ASC specks to the cytosol at subsequent timepoints (Kerur et al., 2011). However, ASC did not form a single large speck in the nucleus or cytosol of this cell type, but rather a speckled pattern with multiple puncta. Another study in zebrafish demonstrated that nuclear ASC speck formation can happen in keratinocytes in vivo (Kuri et al., 2017). In this case, nuclear ASC speck formation in keratinocytes was associated with cell death, which we didn’t detect in primed microglia. Finally, another study linked nuclear ASC speck formation in UVB-exposed keratinocytes to cell death (Smatlik et al., 2021). In this study, nuclear ASC speck formation in human primary keratinocytes was sufficient to trigger Interleukin-1α and -1β release, whereas in human microglia, nuclear ASC speck formation elicited by LPS priming did not lead to Interleukin-1β release (Figure 6). Additionally, the caspase-1 cleaved p20 fragment, a complementary readout of caspase-1 autocatalytic cleavage, could only be detected in mouse microglia treated with LPS and Nigericin, and not in vehicle-treated G2019S microglia, which displayed only nuclear ASC specks (Figure 2B). Thus, as opposed to keratinocytes, nuclear ASC speck formation in microglia is insufficient to trigger caspase-1 activation and cytokine release, but may be an intermediate step within the pathway.

To the best of our knowledge, this is the first report of nuclear ASC speck formation in microglia and myeloid lineage cells. We propose that nuclear ASC speck formation in microglia may indicate a state of inflammasome pre-activation in this cell type.

The pattern of caspase-1 activation in LRRK2 mutant microglia was enhanced compared to Wild Type microglia (Figure 2B), suggesting that ASC specks accumulating within the nuclei in resting microglia could exacerbate caspase-1 activation in response to pro-inflammatory stimuli. Overall, our data highlight regional differences in adaptor protein ASC aggregation propensity as well as genotype-dependent differences in primary microglia. The oligomerization of ASC within the nucleus could represent a pool of ready-to-use inflammasome components that are readily transferred in the cytosol under conditions of stress. This potential novel mechanism, which will deserve future investigation, may accelerate and facilitate NLRP3 activation and the downstream proinflammatory cascade.

The implications of these findings are significant for understanding the pathogenic mechanisms underlying PD, particularly in the context of neuroinflammation. The observed differences in the phenotype of mutant LRRK2-G2019S microglia compared to Wild Type microglia, both in mouse models and human monocyte-derived microglia, highlight the potential of LRRK2 as a therapeutic target for modulating neuroinflammation in PD. In addition, our results suggest that the subcellular location of the ASC speck may be associated with different steps of the inflammasome activation pathway, as it was found only within the cell nucleus during priming and in the cytosol in presence of the trigger signal. Thus, discriminating between nuclear and cytosolic ASC specks may be a more precise method to assess whether microglia is completely activated or just primed, clarifying the interpretation of the results.

In conclusion, the findings of this study provide novel insights into the role of the pathogenic LRRK2 G2019S mutation in microglia and its implications for the pathogenesis of PD. Collectively, these results contribute to the growing body of evidence implicating neuroinflammation in the pathogenesis of PD. The observed dysregulation of the NLRP3 inflammasome pathway in the context of the LRRK2 G2019S mutation provides a mechanistic link between genetic and idiopathic forms of PD, suggesting that similar inflammatory pathways may be involved in both forms of the disease. This has important implications for the development of targeted therapies that can modulate neuroinflammation in PD, potentially slowing or halting disease progression. More research in this area is warranted to fully elucidate the underlying mechanisms and explore the potential of targeting LRRK2-mediated neuroinflammation for the development of disease-modifying therapies for PD.

## List of Abbreviations

APC: Antigen-presenting cell
ASC: Apoptosis-associated Speck-like protein containing a CARD
ATP: Adenosine triphosphate
BSA: Bovine Serum Albumin
BzATP: 2′(3′)-O-(4-Benzoylbenzoyl)adenosine 5′-triphosphate
CARD: Caspase Recruitment Domain
CD11b: Cluster of Differentiation 11b
CSF: Cerebrospinal Fluid
DAMPs: Damage-Associated Molecular Patterns
DMEM: Dulbecco’s Modified Eagle’s Medium
DMSO: Dimethyl Sulfoxide
ELISA: Enzyme-Linked Immunosorbent Assay
FBS: Fetal Bovine Serum
GM-CSF: Granulocyte-Macrophage Colony-Stimulating Factor
hMDMi: Human Monocyte-Derived Microglia-like cells
HRP: Horseradish Peroxidase
IBA-1: Ionized calcium-binding adapter molecule 1
IL-1β: Interleukin-1 beta
IL-34: Interleukin-34
LPS: Lipopolysaccharide
LRRK2: Leucine-Rich Repeat Kinase 2
hMDMi: Human Monocyte-Derived Microglia-like cells
M-CSF: Macrophage Colony-Stimulating Factor
NF-κB: Nuclear Factor kappa-light-chain-enhancer of activated B cells
NLRP3: NOD-, LRR- and Pyrin Domain-Containing Protein 3
PAMPs: Pathogen-Associated Molecular Patterns
PBMCs: Peripheral Blood Mononuclear Cells
PBS: Phosphate Buffered Saline
PD: Parkinson’s Disease
PFA: Paraformaldehyde
PMA: Phorbol 12-myristate 13-acetate
PVDF: Polyvinylidene Fluoride
RCF: Relative Centrifugal Force
RIPA: Radioimmunoprecipitation Assay Buffer
ROS: Reactive Oxygen Species
SDS-PAGE: Sodium Dodecyl Sulfate Polyacrylamide Gel Electrophoresis
SEM: Standard Error of the Mean
SNpc: Substantia Nigra pars compacta
TBS-T: Tris-buffered Saline with Tween 20

## Declarations

## Ethics approval and consent to participate

The housing and handling of the mice used in this study were carried out in accordance with national guidelines. All procedures performed in mice were approved by the Ethical Committee of the University of Padova and the Italian Ministry of Health (license Novello D2784,N,WAT). Human blood samples were collected at the Clinical Investigation Centre (CIC; APHP: ICM); written and informed consent was obtained from all participating subjects. Healthy subjects (controls) were checked for any chronic or acute inflammatory disease, and they did not suffer from any neurological disorder.

## Availability of data and materials

The raw data generated during the current study is available from the corresponding author on reasonable request.

## Author’s contribution

LBa and TB contributed equally to this work. EG, OC, LBa and TB, designed the study. LBa performed the experiments, analyzed and interpreted data on primary mouse microglia. CD collected blood from the LRRK2 patients, isolated PBMCs and set up first differentiation conditions into microglia-like cells. TB performed the experiments, analyzed and interpreted data on patient hMDMi. LI and HW supervised and assisted in primary microglia and hMDMi experiments, respectively. JCC and SS supervised the studies involving patient samples. EG, OC and SN contributed to data analysis and interpretation. LBa and TB wrote the manuscript. SN, EG, OC and LBu have contributed to manuscript editing and optimization. All authors have read and approved the final manuscript.

## Acknowledgments

We express our sincere gratitude to the DeBio Imaging Facility at the University of Padua for sharing their expertise and for their technical support, as well as to the members of the Laboratory of Brain Physiopathology for their critical review and valuable feedback. We are grateful to Adeline Muscat for technical support with the isolation of PBMCs and the generation of MDMi. We acknowledge Fondation Yolande Calvet and Fondation pour la Recherche Médicale for granting a research fellowship to Cédric Dusanter. Finally, we are grateful to Graziella Mangone, Sara Sambin, and all the clinicians contributing to the NS-Park Consortium.

---

**Supplementary Figure 1.**
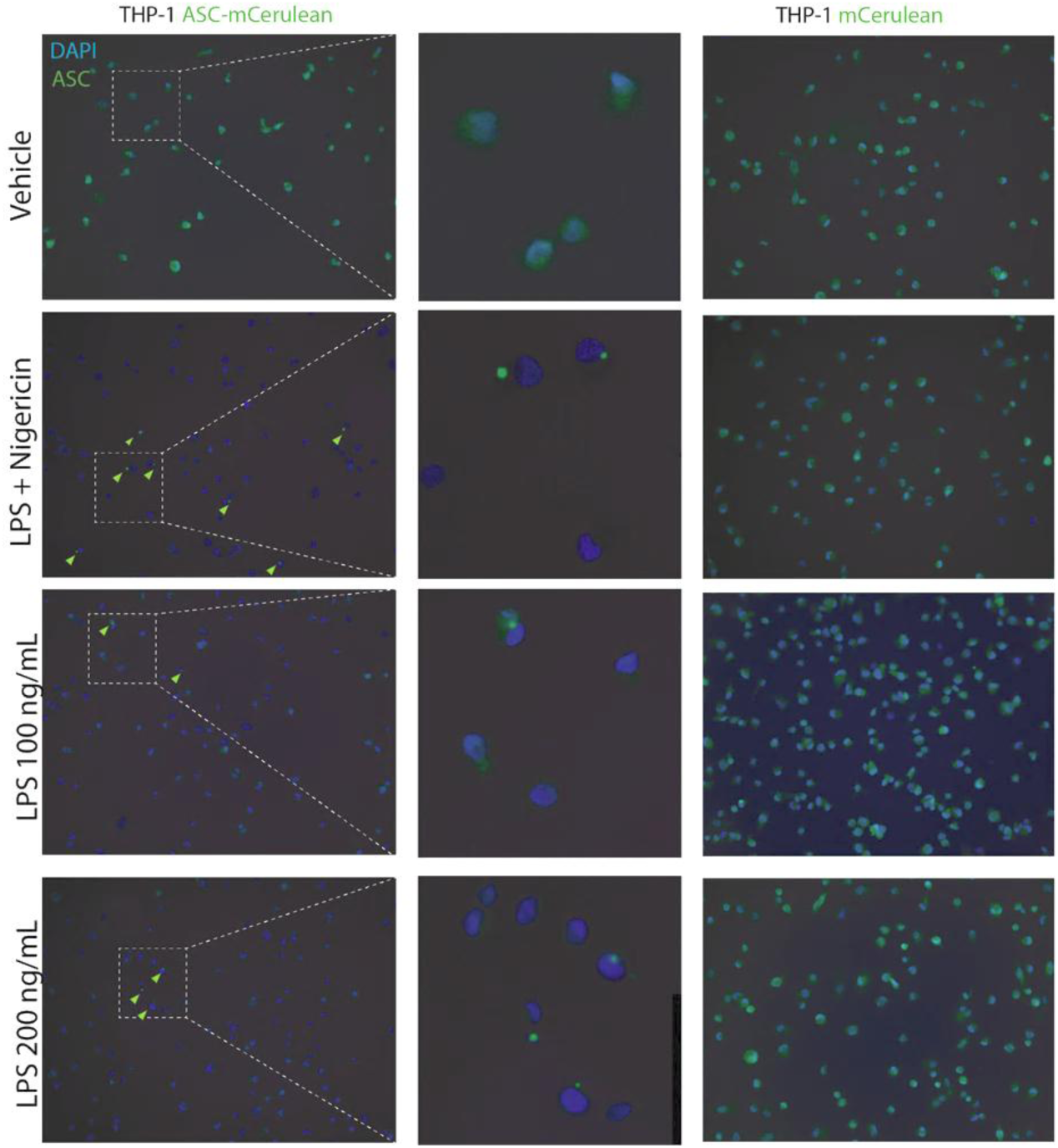
Absence of nuclear ASC specks in THP-1 macrophages. THP-1 differentiated macrophages stably expressing ASC-mCerulean were treated with either LPS (100 ng/mL) + Nigericin or LPS alone (100 or 200 ng/mL). In all treated conditions ASC specks were detected either in the cytosol or perinuclear area.

**File name:** Additional file 1

**File format:** MP4

**Title of data:** Nuclear ASC speck in G2019S microglia

**Description of data:** 3D clip of a nuclear ASC speck detected in LRRK2-G2019S primary mouse microglia.

